# Noise-correlation is Modulated by Reward Expectation in the Primary Motor Cortex Bilaterally During Manual and Observational Tasks in Primates

**DOI:** 10.1101/2020.03.09.984591

**Authors:** Brittany Moore, Sheng Khang, Joseph Thachil Francis

**Affiliations:** Department of Biomedical Engineering, Cullen College of Engineering, The University of Houston, Houston, TX 77204, United States; Department of Electrical and Computer Engineering, Cullen College of Engineering, The University of Houston, Houston, TX 77204, United States

**Keywords:** Motor Cortex, Reward, Reinforcement Learning, Noise-correlation, Reward Prediction, Brain Computer Interface

## Abstract

Reward modulation is represented in the motor cortex (M1) and could be used to implement more accurate decoding models to improve brain computer interfaces (BCIs) (Zhao et al. 2018). Analyzing trial-to-trial noise-correlations between neural units in the presence of rewarding (R) and non-rewarding (NR) stimuli adds to our understanding of cortical network dynamics. We utilized Pearson’s correlation coefficient to measure shared variability between simultaneously recorded units (32 – 112) and found significantly higher noise-correlation and positive correlation between the populations’ signal- and noise-correlation during NR trials as compared to R trials. This pattern is evident in data from two non-human primates (NHPs) during single-target center out reaching tasks, both manual and action observation versions. We conducted mean matched noise-correlation analysis in order to decouple known interactions between event triggered firing rate changes and neural correlations. Isolated reward discriminatory units demonstrated stronger correlational changes than units unresponsive to reward firing rate modulation, however the qualitative response was similar, indicating correlational changes within the network as a whole can serve as another information channel to be exploited by BCIs that track the underlying cortical state, such as reward expectation, or attentional modulation. Reward expectation and attention in return can be utilized with reinforcement learning towards autonomous BCI updating.

## Introduction

The neural representation of reward has been traced to deep brain structures including the substantia nigra pars compacta and ventral tegmental area (Schultz, Dayan, and Montague 1997). Reward associated signals have also been shown to occur in multiple cortical areas including the orbitofrontal and sensorimotor cortices (Tremblay and Schultz 1999; Ramkumar et al. 2016; Ramakrishnan et al. 2017; Zhao et al. 2018; An, Yadav, Badri Ahmadi, et al. 2018; McNiel, Choi, et al. 2016a; Marsh et al. 2015). Previous studies have linked cortical responses to dopaminergic pathways, which can explain cortical responses on multiple time scales (Kunori, Kajiwara, and Takashima 2014) and indicates the complex interactions that exist between deep structure reward signaling and M1 units. For example, dopamine has proven necessary for learning in M1 (Molina-Luna et al. 2009) and has a significant effect on long term potentiation (LTP) in the motor cortex, for review see (Francis and Song 2011).

Studies of the motor cortex reveal a complex dynamical system (Shenoy, Sahani, and Churchland 2013). In addition to kinematic, directional (Georgopoulos et al. 1982), and force tuning information (Chhatbar and Francis 2013; Graham et al. 2003), M1 neurons encode value and demonstrate reward modulation (Marsh et al. 2015; An et al. 2019; An, Yadav, Ahmadi, et al. 2018; Ramkumar et al. 2016; Ramakrishnan et al. 2017; Zhao et al. 2018), as do regions associated with motor output (Platt and Glimcher 1999; Zhao et al. 2018; Musallam et al. 2004; Tanaka et al. 2004; Campos et al. 2005; Shuler and Bear 2006; Louie, Grattan, and Glimcher 2011; Toda et al. 2012). Models indicate that signals encoding value and reward expectation in M1 play a significant role in the reinforcement learning process (Tarigoppula et al. 2018; Dura-Bernal et al. 2015). This information is encoded during active movement, as well as during action observation and motor planning which further implies an interconnection between reward, learning, and expectation (Tkach, Reimer, and Hatsopoulos 2007; Dushanova and Donoghue 2010; Marsh et al. 2015).

Many of the single and multi-unit activity studies on reward modulation in M1 have utilized firing rate, with some exceptions (An et al. 2019; Marsh et al. 2015; An et al. 2018), where the local field potential was studied. However, the correlation between individual units can also provide insight into how populations of neurons change with the task at hand (Maynard et al. 1999; Lee et al. 1998). There is notable variability in individual unit responses to repeated stimuli (Tolhurst, Movshon, and Dean 1983), and this variation is often shared between neurons (Shadlen and Newsome 1998). Research has shown that the correlation between trial-to-trial fluctuations has a significant impact on coding efficiency, specifically in relation to attention, learning, and behavior (Zohary, Shadlen, and Newsome 1994; Abbott and Dayan 1999; Nirenberg and Latham 2003; Averbeck and Lee 2006; Cohen and Newsome 2008).

Changes in correlation occur even in the absence of changes in firing rate, suggesting that correlation dynamics are a critical part of the neural code (Vaadia et al. 1995). Correlation reveals neural population interactions that contribute to our understanding of network architecture. Analysis of changes in correlation in different contexts indicates increased correlation between similarly tuned ensembles and local circuits, affecting learning-related plasticity (Komiyama et al. 2010). In addition, correlation has proven useful in understanding potential connectivity across regions and populations (Reid and Alonso 1995).

In studying the correlation of M1 units in the presence of rewarding and non-rewarding stimuli, there are several factors to consider. Previous work has demonstrated correlation changes related to spatial attention, learning, and other aspects of the internal neural state. These factors are likely to coexist in a reward task experimental paradigm with conditioned visual cues and may also be driving factors in our results. However, fluctuations in aspects of cortical state may occur on distinct and significantly different timescales, such as attentional shifts vs. circadian related changes. Second, correlation can be significantly affected by the strength of the stimulus response (Ecker et al. 2010), which was shown to contribute to 33% of the across-study variance reported in V1 (Cohen and Kohn 2011). Lastly, the composition of the population plays a role in the relationship between correlation and coding efficiency, which means that the results obtained may be interpreted differently depending on the ensemble of units with predictable variations across subpopulations. For example, while increasing correlation has proven detrimental in homogenous populations and similarly tuned neurons, it may increase information carrying capacity in heterogeneous populations and differently tuned units (Chelaru and Dragoi 2008; Abbott and Dayan 1999). As our task utilized a single target, we did not probe such relationships with tuning properties (see methods).

The goals of this study were to 1) determine the extent and direction of changes in correlation in response to rewarding and non-rewarding visual cues on M1 neural activity and 2) interpret the role of correlation in improved neural encoding of rewarding vs. non-rewarding trials. For example we found that force M1 tuning curve amplitudes are larger during rewarding trials as compared to non-rewarding (see Fig.4 in (Zhao et al. 2018)). Given the capacity for reward modulation observed in previous studies within the sensorimotor cortices (Marsh et al. 2015; David B. McNiel et al. 2016; D. B. McNiel et al. 2016; An et al. 2019; Ramkumar et al. 2016; Ramakrishnan et al. 2017; Zhao et al. 2018), and the association of decreased correlation with increased encoding accuracy, we predicted that the mean response of the M1 population would show noise-correlation decreases during rewarding trials, and that this decrease would include not only reward modulated units, but the M1 network at large. We recently showed evidence in line with this hypothesis in the interaction between neural spike trains and the underlying local field potential (LFP) in M1 (An et al. 2019) with higher phase-amplitude coupling and spike-field coherence seen during the NR trials as compared to R trials, however we did not report on interactions between single and multi-units, such as changes in the correlational structure between pairs of units as we do here.

**Figure 1.**
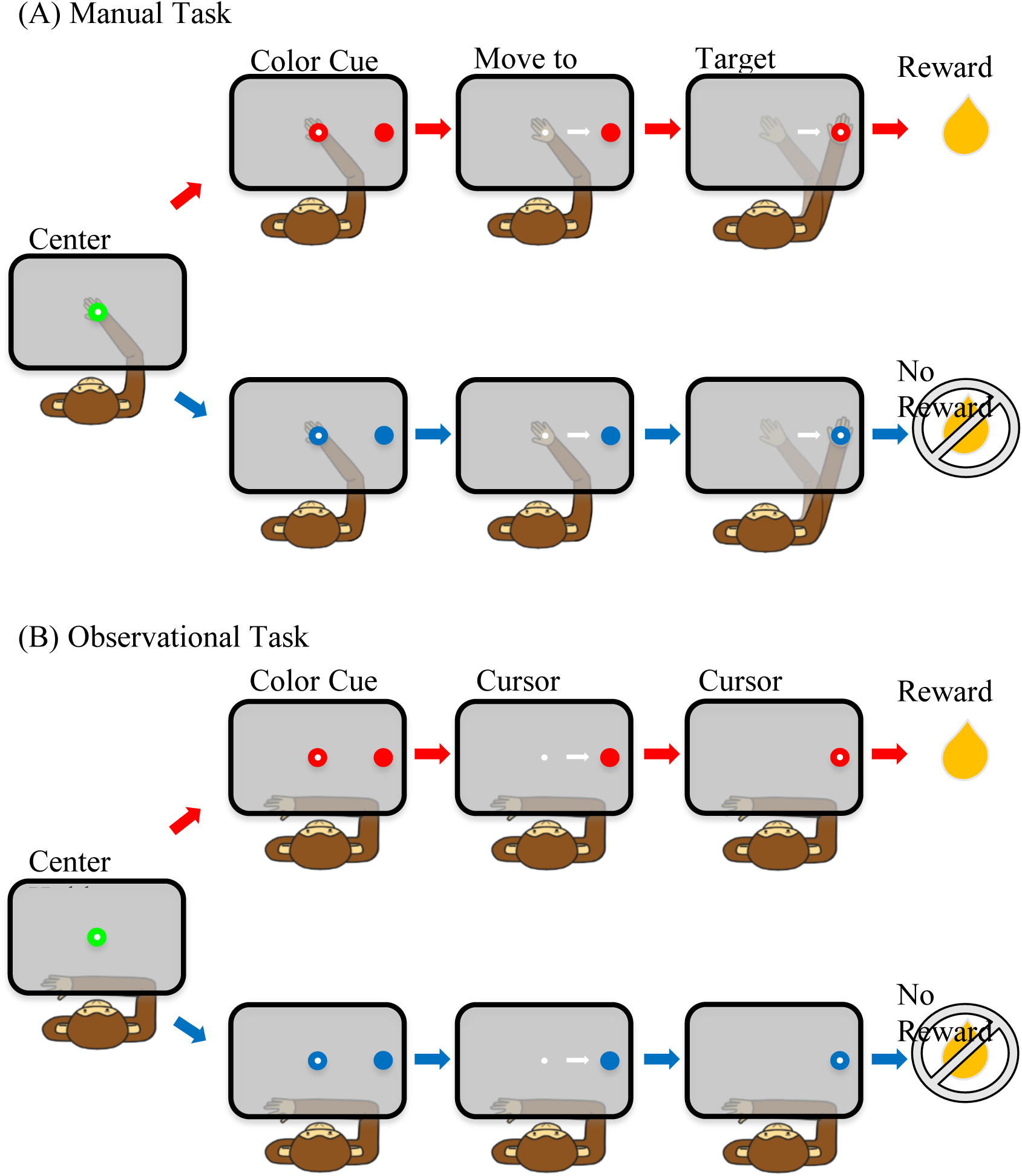
Schematic of the single target reaching task. (A) manual task and (B) observational task.

**Figure 2.**
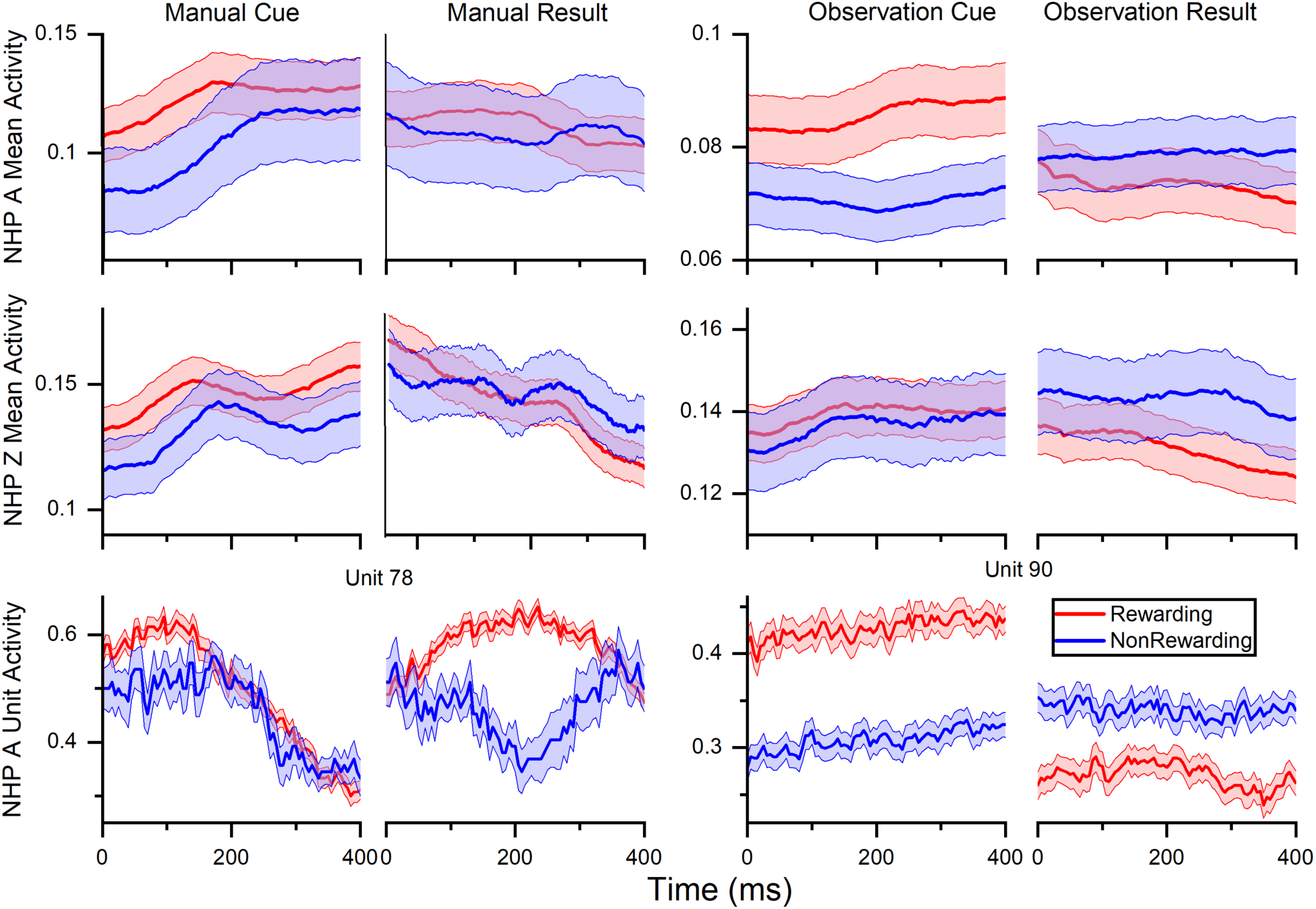
Normalized (min-max) mean firing rate ± SEM for rewarding (red) and non-rewarding (blue) trials averaged across all units and sessions as follows, NHP A manual, N = 80, 80, 112, 80 units across 4 sessions, NHP A observational, N = 91 across all three sessions, NHP Z manual N = 38, 36, 32 units across 3 session and NHP Z observational N = 42, 35, 40 units across 3 sessions.

**Figure 3.**
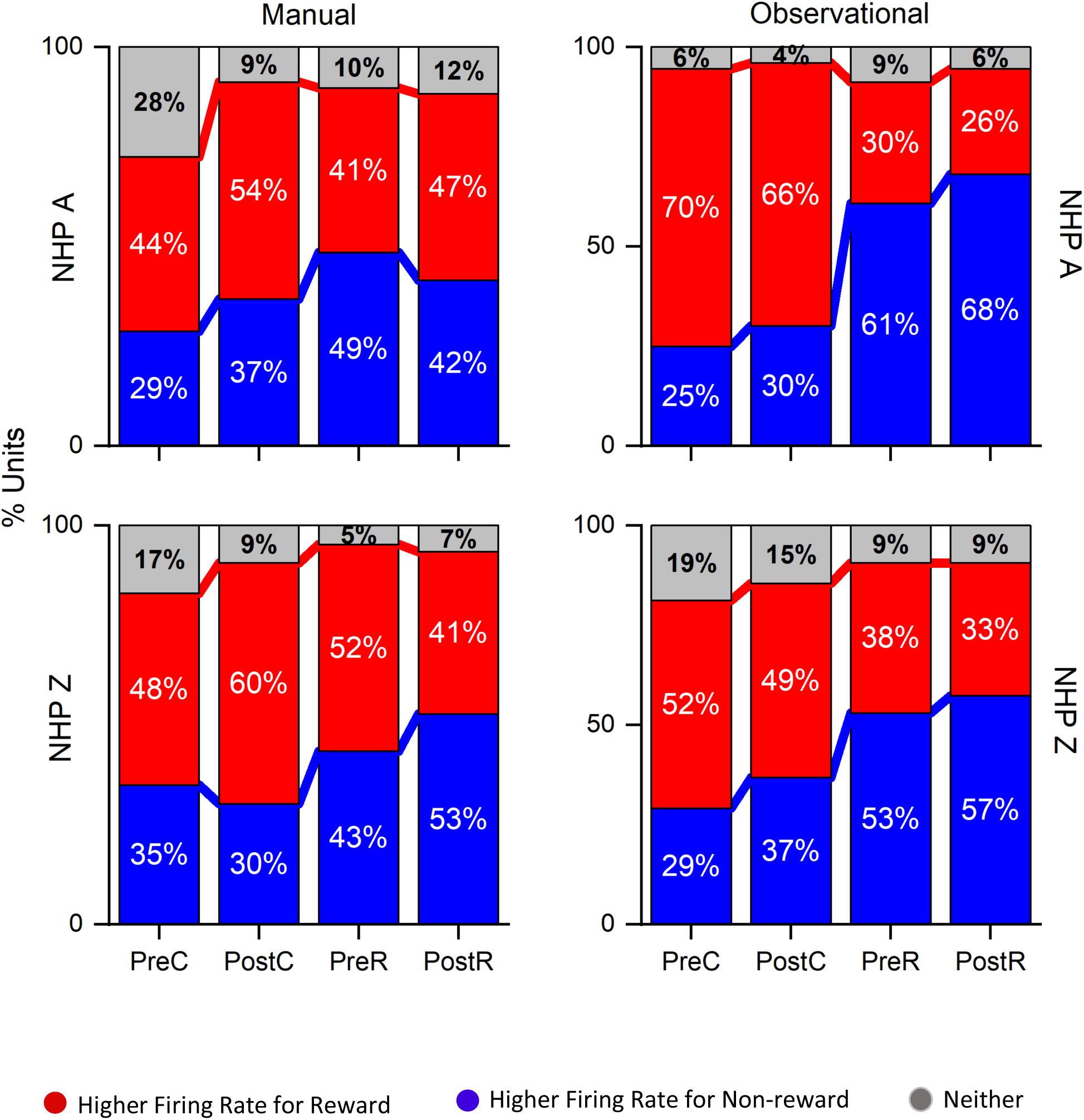
Percentage of units with significantly higher firing rates for rewarding (red) and non-rewarding (blue) trials in each time period, pre-post-cue, pre- and post-result, as well as the percentage of units that were not significantly reward level modulated. The number of units for each NHP and task are as shown in Fig.2 at approximately N = 272 NHP A manual, 273 NHP A observational, 100 NHP Z manual and 117 NHP Z observational, see methods. There were 3 sessions for each NHP and task except for NHP A manual with 4.

**Figure 4.**
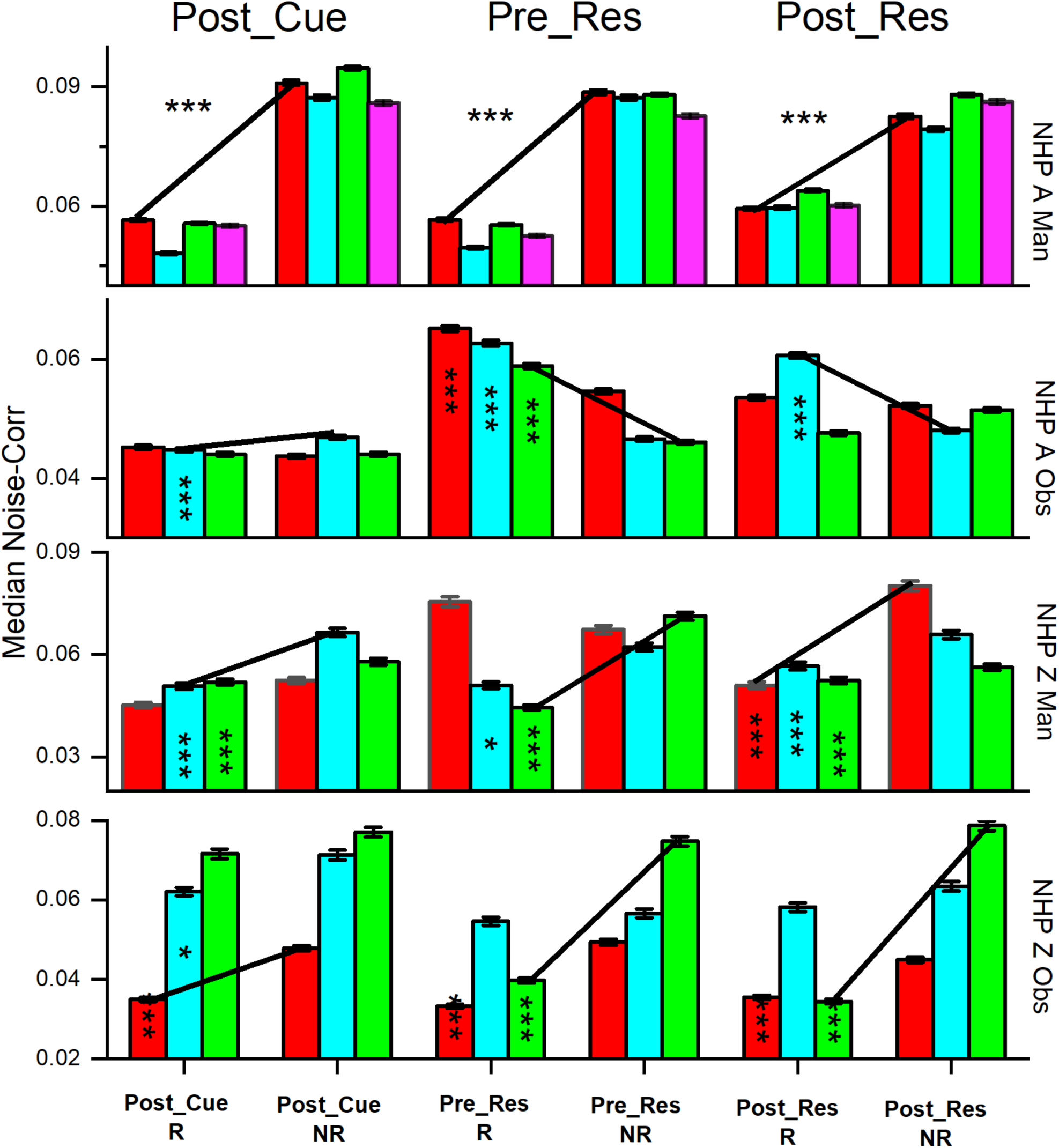
Median absolute value of noise correlations (NC) ± SEM, for rewarding (R) and non-rewarding (NR) trials during the post-cue, pre-result and post-result periods. Significant differences were determined by Wilcoxon rank sum (p < 1 × 10^−10^ and p << 1 × 10^−10^, denoted by *, and *** respectively). All bars deemed significant for the rank sum test were also significant utilizing the Kolmogorov-Smirnov test. Trial type is shown at the bottom and color indicates the data set # for the individual NHPs in order from left to right (red dataset #1, cyan dataset #2 etc.), see methods for more on these sets. The black lines are used to show the trend between the R and NR trial types. We have included the asterisks in the left bars for a pair being compared (R vs. NR).

## Methods

The data used in this study was originally collected and processed as described in (Marsh et al. 2015).

### Surgery

Two bonnet macaques (*Macaca radiata*) were implanted in M1 with a 96-channel microelectrode array (10 × 10 array separated by ∼400 μm, 1.5 mm electrode length, 400 kOhm impedance, ICS-96 connectors, Blackrock Microsystems) using techniques detailed in our previous work (Chhatbar et al., 2010). All surgical procedures were conducted in compliance with guidelines set forth by the National Institutes of Health Guide for the Care and Use of Laboratory Animals and were further approved by the State University of New York Downstate Institutional Animal Care and Use Committee.

Non-human primates (NHPs) were implanted with a headpost (Crist Instrument) three to six months before the electrode implantation. Implantation was performed after animals were trained to complete the manual task (see below) with a success rate of 90%.

#### Data Collection

After 2-3 weeks of recovery, single-unit and multi-unit activity were recorded using multichannel acquisition processor systems (Plexon). The signals were amplified, bandpass filtered from 170 Hz to 8 kHz, and sampled at 40 kHz before the waveforms were sorted using principal component methods (Marsh et al. 2015). There was no segregation between single (SU) and multi-units (MU) for the purposes of our analysis, as the use case is towards BCI applications where both SU and MU have been shown useful in providing information. For NHP A, data was taken from the contralateral M1 and from the ipsilateral M1 for NHP Z with respect to the arm used in the reaching task.

For NHP A manual task, there were 4 same-day recording sessions included in this analysis with 80, 80, 80 and 112 units detected after offline sorting from sessions 1, 2, 3 and 4 respectively. For NHP A’s observation task, there were 3 same-day recording sessions included with 91 units recorded in each using the same MAP sort file (Plexon) indicating these may be the same units (SU and MU). Number of trials for NHP A were, N = 190 R and 61 NR for manual and 469 for both R and NR observational trials. The data collected from NHP A observational task was a perfectly predictable sequence of R trials followed by NR and repeating. This structure allowed the NHP to learn the trial value sequence as shown in (Tarigoppula et al. 2018) with this same data. For Monkey Z manual task, there were 3 different-day recording sessions used for this analysis with 38, 36, and 32 units detected after offline sorting (Offline Sorter from Plexon). For the observation task, there were also 3 different-day recording sessions included with 42, 45, and 40 units recorded in each. Number of trials for NHP Z were, N= 326 R and 159 NR for manual, and 519 R and 258 NR for observational data. It should be noted that the total number of units recorded from was most likely less than the full numbers above, especially for the same day sessions of NHP A as many of the units may have been the same between sessions, however, for this analysis, we did not separate SU and MU data explicitly.

### Experimental Task

The non-human primates (NHP) performed two tasks, manual and observational, that were used for analysis. First, the animals were trained to perform a single target center-out reaching task using their right arm inside a KINARM exoskeleton (BKIN Technologies). For the manual task, a right-hand movement was made from a center target to a peripheral target (0.8 cm radius) located 5 cm to the right (Figure 1). A cursor provided visual feedback of the hand position. The NHP initiated each trial by holding on the neutral (green) center target for 325 ms, followed by a variable color cue period (100-300 ms) where the colored peripheral target appeared as red of blue to indicate a rewarding or non-rewarding trials, respectively. At the same time, the center target turned to the same color as the peripheral target. After a required hold period of 325-400 ms, a GO, marked by the disappearance of the center cue, indicated the NHP could move to the peripheral target where it had to hold for 325 ms (Figure 1A). At that time the animal would receive a liquid reward (R) or no reward (NR), depending on the trial type. If the NHP failed to correctly finish a trial, the same trial type would be repeated. For the manual task, the trial types were randomized otherwise. For the observation task, the trials were presented in a sequenced pattern alternating between R and NR trial types for NHP A. For NHP Z the observational data was biased with a 2 to 1 ratio in favor of R trials, but otherwise randomized.

For the observational task (OT), the NHPs observed the hand feedback cursor as it automatically moved at a constant speed from the center to the peripheral target (Figure 1B). The KINARM was locked into place and the NHP’s left arm was restrained to prevent reaching movements. Eye-tracking was used to ensure that the NHP was looking at the computer projection in the horizontal plane on a semi-transparent mirror where the tasks were displayed during the OT trials.

For each trial, there were 4 defined task phases or period for analysis: pre cue (500 ms), post cue (500 ms), pre reward (500 ms), and post reward (500 ms).

## Data Analysis

### Firing Rate

To determine differences between rewarding and non-rewarding trials, firing rates were calculated using overlapping bins of 100 ms moving in increments of 5 ms. For each unit, *n*, the firing rate, 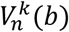 of each bin, *b*, of each trial, *k*, was determined by

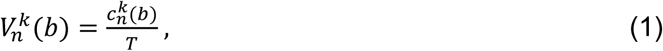

where 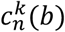 is the number of spike counts per bin and *T* is the length of the bin in ms. For each unit, the firing rate was normalized across the entire session using min-max normalization,

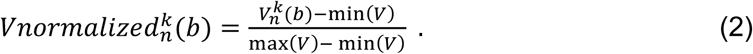

The normalized firing rate, 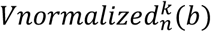, is distributed between 0 and 1. For certain analyses the firing rate data was not normalized, such as when looking at the influence of rate on noise-correlation described below.

### Noise Correlation

Noise-correlation refers to the trial-to-trial variations that are shared between pairs of units (Cohen and Kohn 2011). Noise-correlation is equivalent to the Pearson crosscorrelation of the trial-by-trial firing rate variation of two units about their respective bin-wise means (Bair, Zohary, and Newsome 2001; Kohn and Smith 2005). The trial-by-trial firing rate variation was calculated for each unit, *n*, by

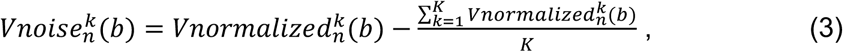

where *K* was the total number of trials. The Pearson coefficient of *Vnoise* was found by

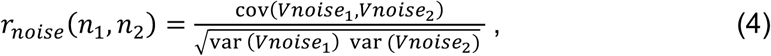

and was calculated using the MATLAB function “corr”. Correlation was calculated separately for each trial period, and all trials of the same trial type were concatenated to find a single coefficient per unit pair per task phase.

### Rate vs. Noise-correlation

The effect of firing rate on noise-correlation was quantified by analyzing the relationship between the arithmetic mean of the units raw activity determined for R and NR data separately. The noise-correlation coefficients during the post-cue response period of all unit pairs was determined for R and NR data separately. Next, we determined which unit pairs from all units had mean absolute differences in their mean post-cue period rate less than 0.8 Hz, which was chosen such that for each data set there was data within each bin. We then took these pairs of units from the R and NR datasets and formed 10 bins, with each bin containing data with a mean individual unit rate within the 0.8 Hz bin. We then plotted this mean rate (x-axis) against the mean of the absolute value of the noise-correlation between the unit pairs within that rate bin. We ran a Wilcoxon Signed Rank test (signrank test in MATLAB set with ‘tail’, ‘left’) on the data as well as a Kolmogorov-Smirnov test (kstest2 in MATLAB). We focused on the post-cue response period, which was consistently different between R and NR trials for all tasks and NHPs for the median noise correlations (Fig.4). Unit pairs from all sessions for a given NHP and task type were combined for this analysis (Fig.5). For individual session median noise correlations see Fig.4.

**Figure 5.**
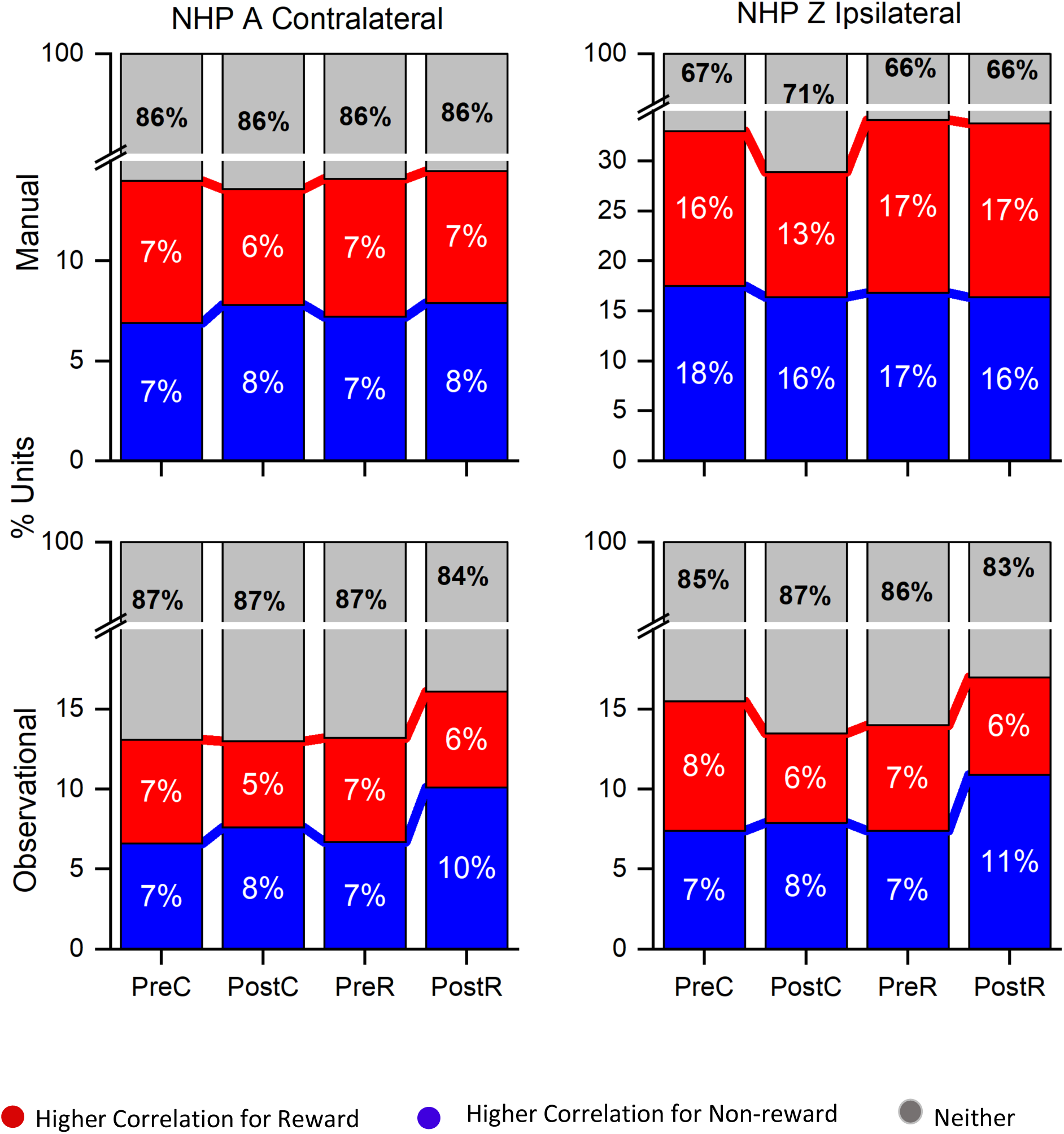
Percent of unit pairs that have higher noise-correlation for rewarding (red), non-rewarding (blue), or neither (gray) trial type for both NHPs and tasks as labeled during the pre- and post-cue, pre- and post-result time periods. Note the break in the y-axis.

### Relationship between Signal and Noise correlation

We conducted analysis of covariance (ANCOVA) between the signal- and the noise-correlation for all unit pairs recorded during R and NR trials separately. We utilized the MATLAB function aoctool for this analysis with the model set to ‘separate lines’, such that there were no added constraints to the model. We have plotted for each unit pair the signal correlation (x-axis), which is simply the crosscorrelation between the mean unit responses (PSTH, post-stimulus-time-histogram) during the given task period, against the noise-correlation for the unit pair. The aoctool outputs statistical confidence on the model parameter values as well as an ANOVA table on the signal vs. noise, the trial type of R vs. NR, and an interaction of Signal-correlation * Reward level vs. noise-correlation, and we have included the ANOVA outputs on the related figure, or accompanying text (Fig.6).

**Figure 6.**
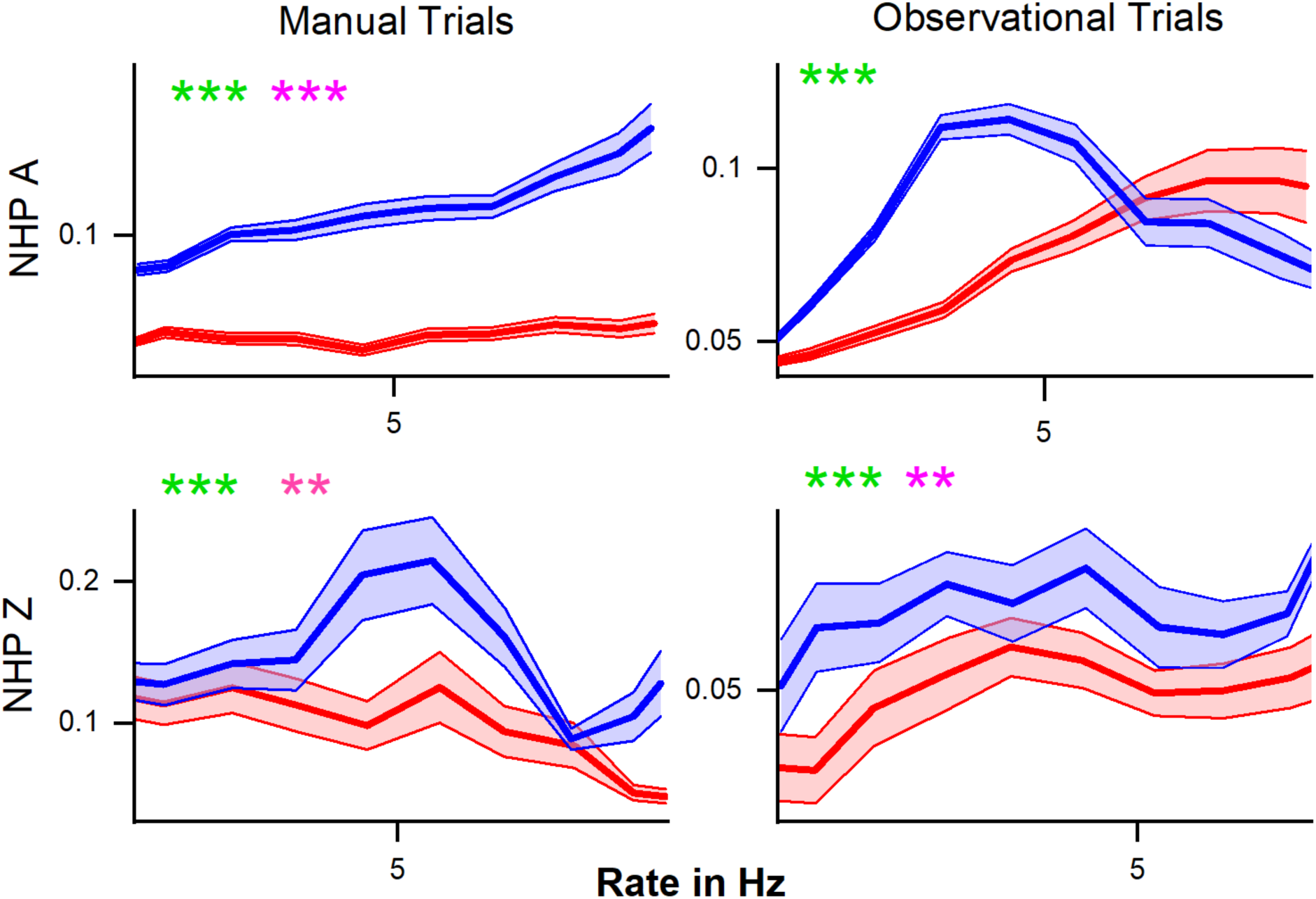
Mean noise-correlation coefficient (y-axis) plotted against post-cue units mean rate for unit pairs with similar mean rates (<0.8 Hz difference). Rewarding (red) and non-rewarding (blue) trials are plotted ± SEM for all unit pairs over all data sets for that NHP and task type. For each subplot we have included statistical results from both the Kolmogorov-Smirnov test (green) as well as the Wilcoxon rank sum test (pink) with * p< .05, ** p<.01, and *** p< .001, between rewarding and non-rewarding trials.

### Identifying Reward Modulated Subpopulations

Selecting reward discriminatory units was achieved using the Pearson correlation between reward value on a given trial and unit firing rate, where R was assigned a value of 1 and NR was assigned a value of 0. We arbitrarily used the top 20% of units with the highest correlation coefficients, showing strong positive correlation with reward, were classified as “High Positive” units. The bottom 20% of units with the lowest correlation coefficients, showing strong negative correlation with reward, were classified as “High Negative” units. Lastly, the middle 20% of units within the middle of the distribution of absolute value correlation coefficients, were classified as “middle-responsive” units. These subpopulations were determined for each time period separately, which means they may consist of different units during each period. This analysis was carried out in order to determine if all three types of units would still show modulation in their noise-correlation and if these populations would differ towards finding the best information channels on cortical state related to reward expectation, reward associated motivational salience or attention.

## Results

We present our results addressing our main hypothesis that reward expectation influences trial-to-trial noise-correlation, such that this measure could be used to track the cortical state of expecting reward vs. not expecting reward, which we aim to use towards autonomously updating brain computer interfaces. Toward this aim we recorded from neural ensembles simultaneously (30 – 112 units) in contralateral or ipsilateral M1 in NHPs performing a single target reaching task, or observing such a task, as seen in Fig.1. To present a fuller story we first show raw data that indicates reward expectation modulates M1 rates as we have previously described (Marsh et al. 2015; Tarigoppula et al. 2018; An et al. 2019). We start our results with raw PSTHs between R and NR trials (Fig.2), moving onto inquiry about the % of units that modulate their rate to reward-level cueing (Fig.3), after which we show the mean noise-correlation for each dataset for R and NR trials (Fig.4). We then looked at how firing rate influences noise-correlation using mean matched methods (Fig.5) and then show correlational structure between the noise-correlation and the signal correlation (Fig.6). Finally, we look more closely at how subpopulations of units noise-correlation is modulated by reward expectation, such as units which have increased rate modulation due to reward, or decreased rate, or no reward modulation shown in rate, as well as how high positive correlated units are modulated by reward cueing vs. highly negatively correlated units.

In figure 2 we have plotted the average min-max normalized mean firing rate by trial type, rewarding (R, red) and non-rewarding (NR, blue), across all units for all sessions. NHPs (A, contralateral, and Z, ipsilateral) and task types (manual and observational). These results align with previous work showing increased firing rate in response to a preferred stimulus and a higher overall rate for rewarding stimuli (Marsh et al. 2015; Tarigoppula et al. 2018a), and utilized some of the same datasets used here from the same NHPs. This trend can be seen in both the manual and observation tasks, indicating that reward modulation occurs in the motor cortex while performing a reaching movement and during passive observation as expected (An et al. 2019; Marsh et al. 2015). In addition, there are differences between the manual and observational tasks consistent with previous research, where neural responses to observation tend to be weaker than during the manual version of the task. We have plotted two example units for NHP A manual and observational tasks in the bottom row of Fig.2. NHP A’s observational data is separated between R and NR even at the start of the pre-cue period. This separability can be attributed to the predictable nature of the trial value sequence for NHP A’s observational task, which had a repeating sequence of R followed by NR trials, in comparison to the more randomized trial sequence in the manual task (see methods). As NHP A learned the sequence of R-NR the difference in firing rate can be seen before the cue onset as shown in (Tarigoppula et al. 2018) for these data sets. Please note results on the above datasets, and related datasets, for differences in firing rate, duration of trials, and EMG have been presented previously by us showing the expected results, such as slight increased trial duration during manual tasks for NR trials as compared to R trials (Marsh et al. 2015; An et al. 2019; Tarigoppula et al. 2018; Hessburg, Zhao, and Francis 2019). However, the observational tasks have no such differences, and as seen in our results still show clear differences due to reward level cuing.

Figure 3 illustrates the percentage of units in each time period that was modulate by cued reward level. In each graph, red indicates a higher firing rate for R-trials, blue indicates a higher firing rate for NR-trials, and grey indicates no significant difference between the two trial types (Wilcoxon rank sum, p < 0.05). For both NHPs, most of the units modulate for reward when firing rate is compared in each time period. There are a larger percentage of units with significantly higher firing rates for rewarding trials in the post-cue period, and this percentage decreases as the trial phases move forward (Figure 3). During the pre-result and post-result periods of the observational task, both NHPs show a greater percentage of units with increased firing rate in NR trials as shown previously (Tarigoppula et al. 2018).

The differences in firing rate between R and NR trials confirms that reward modulation occurs in the motor cortex during these single target center-out reaching tasks (Marsh et al. 2015), which has also been shown in multi-target tasks (Zhao et al. 2018; Tarigoppula et al. 2018; Ramakrishnan et al. 2017; Ramkumar et al. 2016). This metric alone, however, does not provide a complete perspective on the dynamics of these populations. Further analyzing the interactions between these units is critical to understanding more broadly the cortical state.

### Noise Correlation

The observed noise-correlation (NC) includes both positive and negative coefficients which results in a mean NC close to zero, similar to the range given by Ecker et al., of 0.01 to 0.03 (Ecker et al. 2010). Figure 4 displays the absolute value of the unit pair NC for all unit pairs. The results show NC for R and NR trials, with some variation across time periods, task, and subject. During the manual and observational tasks, as hypothesized, both subjects showed significantly higher NC for NR trials during the post-cue period. During the pre- and post-result time period, both NHPs for the manual task show the same relationship of higher NC for NR trials and so did NHP Z for the observational task. However, NHP A, which had the predictable trial value sequence for the observational task, showed the opposite relationship during the pre- and post-result time periods as seen in the second row of Fig.4.

### Percent of Population

Considering all unit pair noise-correlation coefficients, the significance of the difference between R and NR NC was determined by Wilcoxon rank sum (p-value < 0.05) for each unit pair. Unit pairs were classified as either having a higher NC during rewarding (red) or during non-rewarding trials (blue) (Figure 5). NHP A showed a slightly greater percentage of unit-pairs with higher NR NC in most time periods. A larger percentage of unit pairs do not demonstrate a significant difference between R and NR trials for their NC (gray), suggesting that NC differences on the Unit level are not strong and the population as a whole carries this information (compare Figs.4 and 5). NHP A observation, NHP Z manual, and NHP Z observation data all follow the similar trend with slightly more unit pairs showing higher NC for NR trials (Figure 5). The post-cue period consistently showed a slightly higher percentage of unit pairs with increased NC for NR trials across all NHPs and tasks.

### Effect of firing rate

Previous work indicated that an increase in firing rate causes an increase in NC (de la Rocha et al., 2007). Both NHP A and Z demonstrate higher NC for NR trials regardless of the relationship between post-cue induced firing rate changes and NC. The relationship between mean NC and mean matched firing rates differed between the trial types and NHPs (Fig.6). In NHP A, NC showed a linear increase with firing rate during the manual task for NR trials. However, for the observation task, there was an inverse effect on NR trials that resulted in a sharp decrease in NC at higher firing rates, where data from rewarding trials showed a more linear increase in NC with firing rate (Fig.6). For NHP Z, there was a significant tendency for non-rewarding trials to have higher NC for all rates as seen in Fig.6. However, the relationship between rate and NC was not simply linear, as also seen for NHP A observational data. These relationships remained whether we used the geometric mean (data not shown) or the arithmetic mean. In Fig.6 green asterisks are used to report results from KS-tests, while the pink asterisks are results from Wilcoxon signed rank tests. In Fig.6 we focus on the lower rage of firing rates as R trials, as seen in Fig.2, had higher rates and NR trials would have very few or no rate bins with data in them at those higher rates. However, when using a geometric mean binning method and spanning the full rate space we did not see qualitatively different outcomes (data not shown).

### Signal vs. Noise Correlation

Previous work has shown a relationship between the signal-correlation and the NC, and we wished to determine if the reward expectation level would have an influence on this signal- and noise-correlation relationship. We have plotted the signal correlation for every unit pair for the R and NR trials as red and blue points in scatter plots as seen in Fig. 7, along with linear fits to both data type separately. In both NHPs and tasks there were significant linear model parameters for intercept and slope (R vs. NR) Prob > |T| = 0. In addition, both NHPs for manual data, and NHP Z for observation, had significant differences between signal-correlation*trial-type vs. noise-correlation (Prob > F = 0), however, NHP A’s Observational data did not show this latter relationship significantly, only for trial-type and signal- vs. noise-correlation (Prob > F = 0). Note, there are tens of thousands of points in these scatter plots, so even apparently subtle differences in the lines can be highly significant.

**Figure 7.**
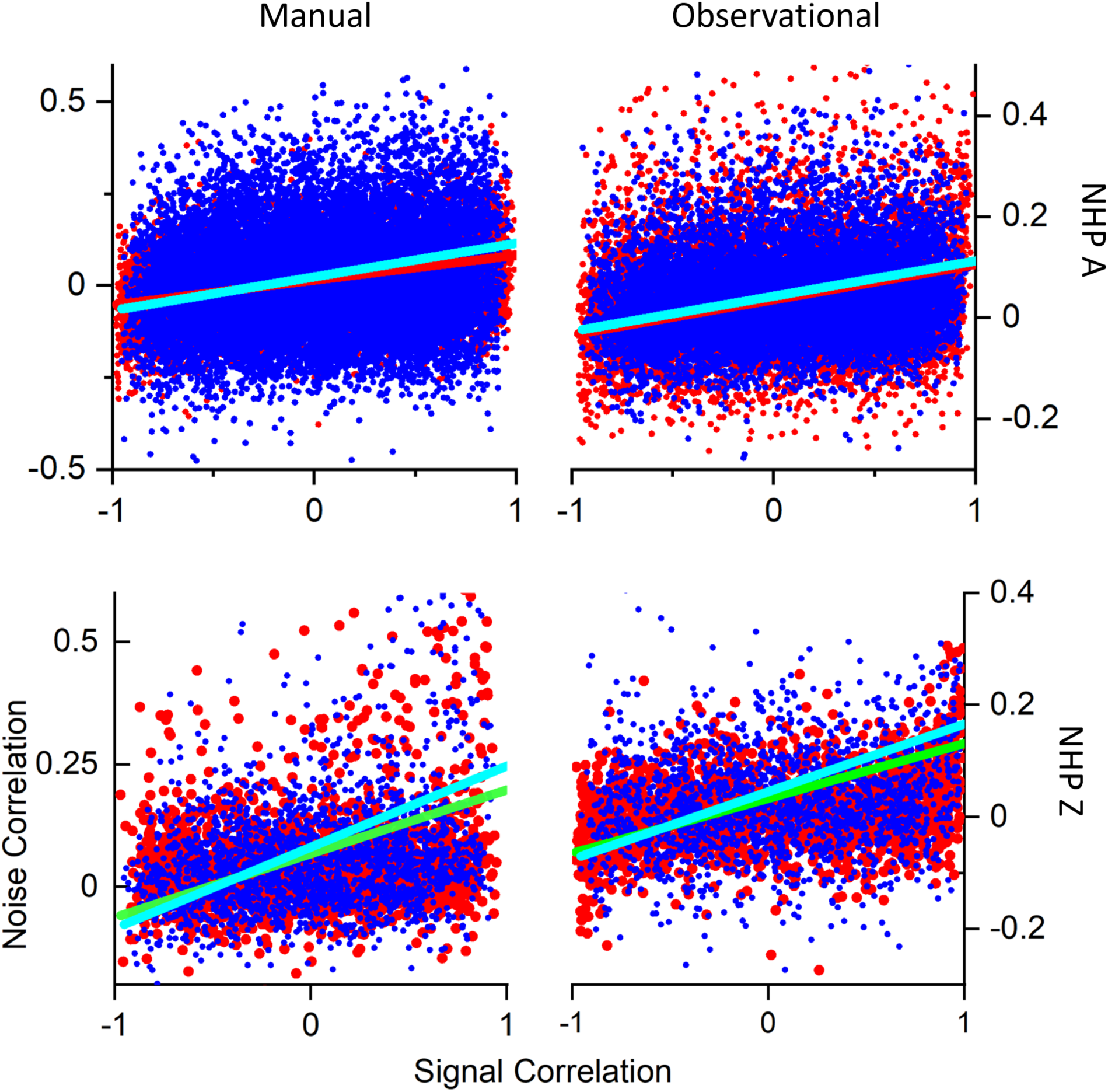
Signal- vs. Noise-correlation. Plotted in red are the rewarding trial data from all datasets for each NHP and task, while blue datapoints are from the non-rewarding trials. Cyan line is a linear fit to the non-rewarding trials and red (NHP A) or green (NHP Z) are linear fits to the rewarding data. The x-axis is signal-correlation while the y-axis is the noise-correlation. All datasets had highly significant (Prob > F = 0 and Prob > |T| =0) linear fits between most of the variables involved, including trial-type, y-intercept and slope of the linear fits. All datasets for NHP Z, and the manual datasets for NHP A also showed highly significant fits for signal-correlation * trial-type vs. noise-correlation (Prob > F = 0). Statistics included ANOVA output for the relationships between signal-correlation, noise-correlation and trial-type. MATLAB aoctool was utilized for this ANCOVA. Number of data sets was (N=3 for all but NHP A Manual N = 4, see methods for trial and unit #s).

### Sign (+/-) of Noise-Correlation and Stability

Figure 8 shows separation of the median positive and negative NC for each NHP and task during the post-cue time period (Fig.8.Top Row). Positive (Pos) and negative (Neg) NC are shown for both R and NR trial types. It is clear that the left two columns of each group of 4 bars, which are for the positive NCs, are higher than the negative two NC bar values seen in the right two columns of each group of 4 bars. Likewise, NC is generally higher during the NR trials as compared to the rewarding trials. Previous studies have related lower information encoding capacity to increases in positive NC, referring to the mean response of a population of neurons (Zohary, Shadlen, and Newsome 1994). Dissecting the distribution of these coefficients by separating positive and negative correlations may explain discrepancies between subjects and task types and reveal additional information about the role of NC (Chelaru and Dragoi 2016). For NHP A manual task, the positive and negative coefficients are both greater in absolute value for NR trials during all time periods within the trials. This aligns with the mean and median NC, which are higher for NR trials (Fig.4 and related text). In general, the other time periods within the task also show greater NC during NR trials for both positive and negative coefficients, and NHP Z observational task displays similar trends to NHP A manual data during all time periods (data not shown).

**Figure 8.**
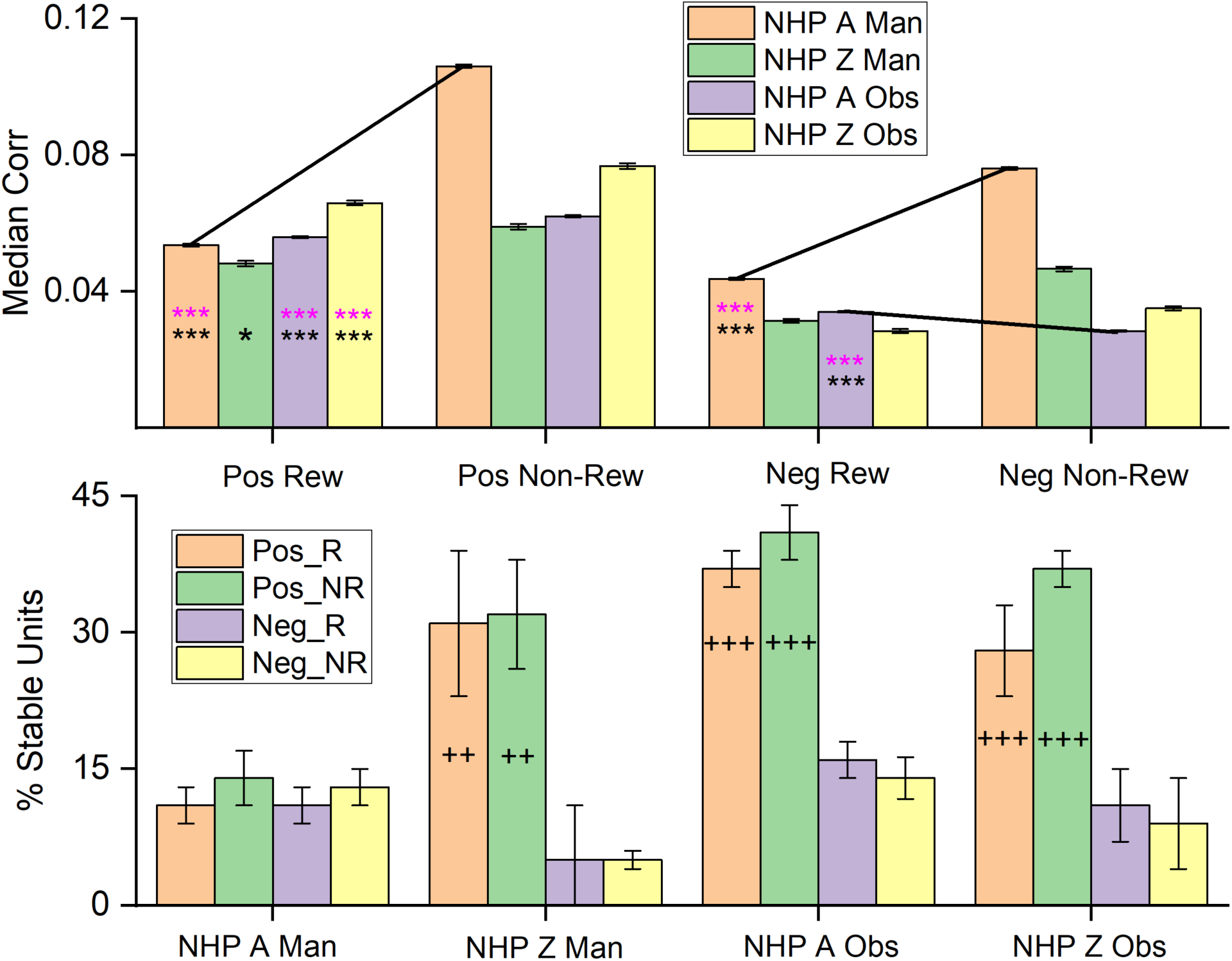
Median noise-correlation during the post-cue period for positive (Pos) and the absolute value of the negative (Neg) noise-correlation coefficients during rewarding (Rew) and non-rewarding (Non-Rew) trials from all data (Top Row, ± SEM, for median, over all data points from all dataset). Significance was determined using KS-test and Wilcoxon signed rank (*, **, ***, where p < 0.05, p < 0.01,p < 0.001 respectively). All comparisons between Pos data sets for rewarding and non-rewarding trials were significant, where significance was determined by Wilcoxon rank sum (p < 0.05) and KS-tests (p<0.05), as were differences between Pos and the absolute value of the Neg trials (all p<<<.001). Standard error of the medians was on the order of 0.0004 - .0008 for the median plots. (Bottom Row) The percentage of units that remain within a single category (Pos_R, Pos_NR, Neg_R, Neg_NR) throughout the task from post-cue, during pre-results and post-result time periods (± SEM over dataset percentages). The Percent stable units were significantly different after post-hoc testing between the groups with + on their bar. The number of + signs indicates how many out of the three datasets showed post-hoc differences between the Pos and Neg subsets for both R and NR trial types (p < .05 post-hoc, MATLAB multcompare with Tukey’s honest significant difference criterion). Note separate keys for top and bottom rows.

The bottom row of Fig.8 shows the stability of the population of units within a given group, such as units with positive NC during R trials, positive NC during NR trials, negative NC during R trials and negative NC during NR trials. In general, one can see that the positive units are more stable in the sense that they don’t flip back and forth between having positive NC during the different task phase, but remain positive in their unit-pair NC, whether during R or NR trial types. This stability was significant for all datasets except for NHP A manual, which was unexpected when compared to the previous results on NHP A manual data set. NHP A was implanted in the M1 contralateral to the reaching arm and therefore during manual trials this cortical region was most in direct control of the reaching movements, and as NC may indicate a decrease in information capacity, this might explain these results. Nevertheless, NHP A manual data still had the same overall trends as the other datasets with R trials being lower than NR trials.

### Relationship Between Reward-rate Modulation and NC

While the combined neural firing rate of all recorded units shows separability between rewarding and non-rewarding trials, there is evidence that this separability can be attributed to select units (Marsh et al. 2015). Analysis of the currently presented data sets revealed a subpopulation with increased firing rate during R trials and another with increased firing rate during NR trials (Marsh et al. 2015), which led directly to our Fig.9 analysis.

While it is apparent that NC differences in response to reward level cueing are present in only a subset of unit pairs (Fig. 5), the defining features of these subpopulations is still unclear. To use noise-correlations as an indicator of reward expectation, selecting subpopulations that are known to modulate for reward through firing rate provides a more direct comparison between these two factors. Figure 9 shows the median absolute value of the NC for R and NR trials for each of the following subpopulations: 1. units with rates that are upmodulated during rewarding trials (High Positive), units with rates that are downmodulated during rewarding trials (High Negative) and units not modulated strongly by reward (Low). NHP A’s contralateral M1 showed the strongest relationship during the manual task with NR trials having higher NC for all subpopulations, and this was also seen for the high positive subpopulation for NHP A’s observational data. Though NHP Z’s ipsilateral cortex showed little of this clear relationship, however, NHP Z’s data still maintained an overall trend of higher magnitude NC during NR trials compared to R trials (Fig.9).

**Figure 9.**
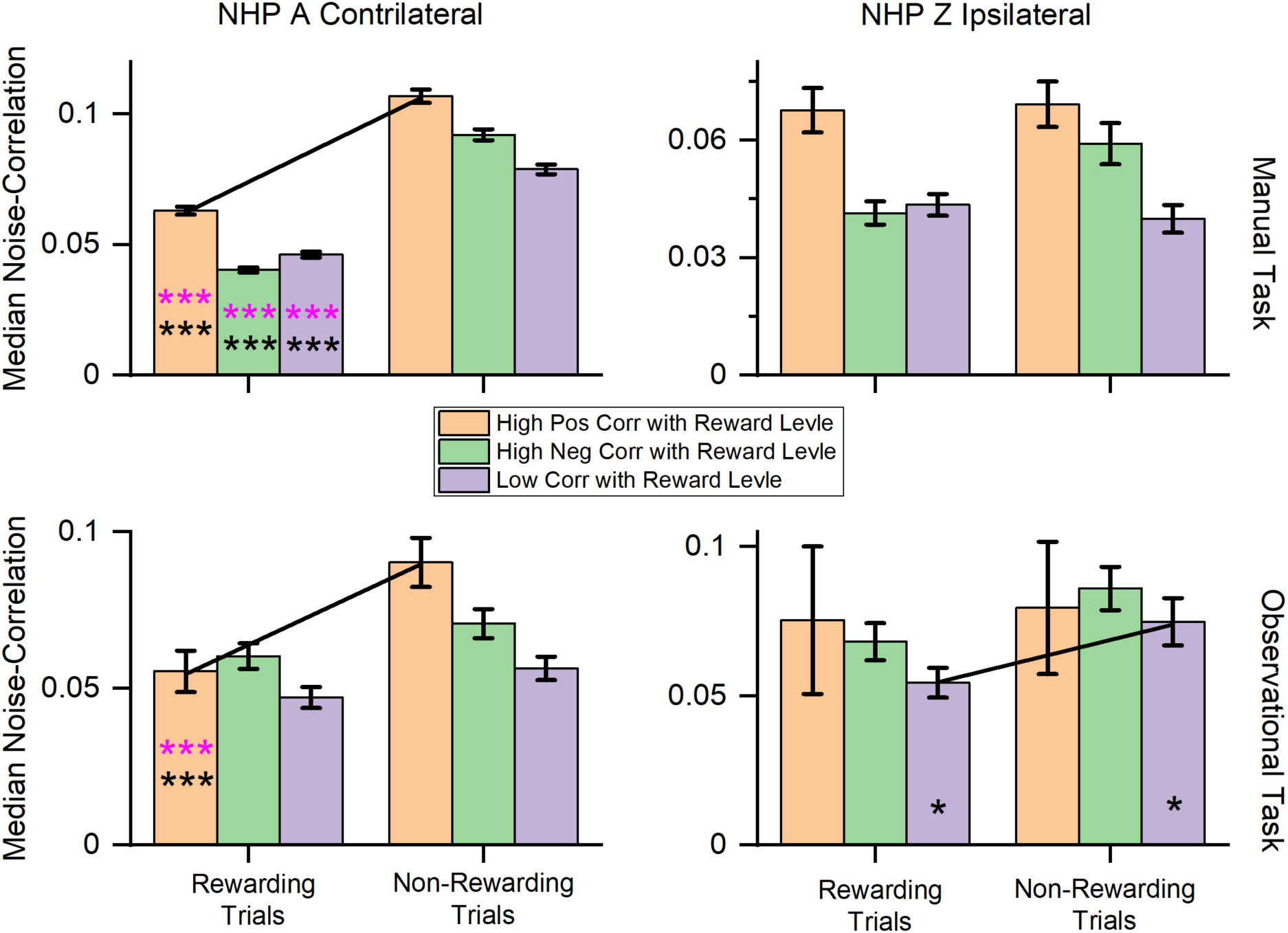
Median correlation coefficient for the High Positive, High Negative and Low-responsive subpopulations each NHP and task as labeled. Black and pink asterisks located on the graph indicate there is a significant difference between the subpopulations during R and NR trials for Wilcoxon rank sum and KStest2 respectively (p < 0.05, and p < 0.001 denoted by * and *** respectively).

## Discussion

The primary goal of this analysis was to determine if significant differences in NC exist between cued R and NR trials, and if so, to what extent might these differences relate with reward modulation of firing rates. As previously described, units with firing rates modulated by reward expectation are present in these data sets (Marsh et al. 2015; Tarigoppula et al. 2018), with one subset of M1 units demonstrating increased firing rates during R trials and one subset showing increased rate during NR trials (Figure 3). Because of the capacity for reward modulation demonstrated by these M1 units during both manual and observation tasks, it was hypothesized that there would be significant differences in NC between R and NR trials. Based on our published local field potential (LFP) work on increased levels of phase-amplitude coupling (PAC) and spike-field coherence (SFC) in the alpha band during NR trials as compared to R trials (An et al. 2019; 2018), we expected to see increases in NC for NR compared to R trials as well. Recent work on these measures (PAC, SFC and NC) in the frontal eye fields has suggested that NC modulations are due to long-range connections via LFP influences (Hassen et al. 2019).

Our presented results follow what would be expect if information capacity within the M1 network was governing the changes in neural rate, and correlational structure. To be clear, we hypothesize the M1 network is carrying the most information about the environment during the task cue period, when the NHP subjects learn the value of the given trial, as well as having information on the cursor position, velocity and the target etc. During this post-cue period both the contralateral and ipsilateral M1s showed an increase in firing rates during R vs. NR trials (Fig.2). Corresponding to this increase in rate was a decrease in the median NC for R trials during the post-cue period (Fig.4), and an increase in NC for the NR trials. A potential factor impacting increased NR NC is firing rate. Previous studies have demonstrated that increases in firing rate are related to increases in correlation (de la Rocha et al. 2007). However, we see the opposite influence of NR on firing rate, that is NR decreases firing rate on average. “Integrate-and-fire” models indicate that output correlation increases in response to increasing firing rate when the input correlation remains unchanged (de la Rocha et al. 2007). This pattern is illustrated by the increase in NC with increasing response firing rates seen in (Fig.6). However, despite the relationship with firing rate, NR trials continue to exhibit higher NC than R trials. The response to changes in rewarding stimuli is similar to the pattern seen in attention, with higher evoked rates leading to greater differences between the two states (Cohen and Maunsell 2009). With increases in response to the value cue, the difference between NR and R NC increased in general, but not all responses were linear, and some decreased after reaching a given rate (Fig.6), and again, NR trials had lower firing rates than R trials. Thus, the increased rate for R trails is accompanied by a decrease in NC, a situation that could allow for more information carrying capacity vs. the NR trials.

In addition to studying the pair-wise correlational structure, we also conducted Principal Component (PC) Analysis (PCA) on our data, finding the same trends seen in Fig.4 between R and NR trial type. Using PCA we asked when the latent dimensionality of the data was lower, during R or NR trials. More of the data’s variance was consistently explained by 20 PCs for NR trials than for R trials, again indicating that there is more correlational structure during the NR trails, leading to a lower dimensional latent space, as compared to the R trials (data not shown). This increased correlation and decreased firing rate during NR trials, shown above, could indicate a cortical state with less information carrying capacity, perhaps as a strategy to conserve energy during low reward expectation moments, which would likely correspond to states of low motivation and decreased attention. By information carrying capacity we mean in the information theoretic sense, where correlation decreases the capacity for a network to transmit information. One can imagine if all the neurons were perfectly correlated it would be similar to having just one neuron, whereas if they are all uncorrelated each neuron could be transmitting independent information at the same time (see, for review or introduction respectively (Magri et al. 2009; Bialek et al. 1996)).

There are several possible explanations underlying the increase in NC during the post-cue period of NR trials including changes in attention, firing rate, motivation, and reward expectation. The first relies on changes in visual attention, where attention may either be a coexisting, or driving factor of correlation, or a separable, but resembling factor that produces similar dynamics to those induced by reward or reward related motivation. Given the predictable nature of the trial reward value sequence (NHP A observation), visual attention could be a confounding factor when discussing the relationship between reward and NC. It is possible that the subjects’ ability to anticipate a lack of reward resulted in a shift to an unattended state during NR trials, even before the trial started. Unattended trials show increases in correlation (Cohen and Maunsell 2009; Herrero et al. 2013; Mitchell, Sundberg, and Reynolds 2009) and could account for the higher correlation coefficients seen in the NR trials. We have shown previously that using a task with multiple levels of reward leads to both linear trends in firing rate with value (Tarigoppula et al. 2018), as well as non-linear gain modulated activity patterns (Hessburg, Zhao, and Francis 2019), making it seem unlikely that all of our results are attributed to attention alone, unless the attention is modulated as one would expect the state-value to be modulated, or the state-motivation. However, further analysis and more directed experiments are needed to convincingly address these questions.

Attention, stimulus contrast (Kohn and Smith 2005), learning (Gu et al. 2011; Jeanne, Sharpee, and Gentner 2013), and global cognitive factors (Ruff and Cohen 2014) have been shown to decrease NC in neurons within the same cortical area. Given the wide variety of factors impacting correlation, the increases during the NR trials could be the result of independent modulatory influence of reward on NC. Previous studies provide insight into the ways in which this difference in correlation between states serves to improve information encoding. High levels of positive correlation can be detrimental to the coding accuracy of a similarly tuned population of neurons (Zohary, Shadlen, and Newsome 1994). Cohen and Maunsell determined that the “modulation of noise-correlation accounts for the majority of the attentional improvement in population sensitivity”, with modulation of firing rate and single unit variabilities accounting for a much smaller portion of the change (Cohen and Maunsell 2009). Similarly, high correlation limits signal-to-noise ratios (SNR), with higher correlations leading to lower saturation of SNR as a function of neuronal pool size (Mitchell, Sundberg, and Reynolds 2009). Much like the trends seen in this research, Mitchell et al., noted that the effects of firing rate on the SNR saturation point does not adequately account for the differences in SNR ratios observed between units with high or low correlation (Mitchell, Sundberg, and Reynolds 2009).

There is evidence that increases in NR trial NC are related to reward modulation. However, the mean correlation does not provide a complete picture of the correlation activity in the population. An additional factor that may be impacting reward modulation is the distribution of positive and negative correlation coefficients. A mean positive correlation reduces the coding capacity of similarly tuned neurons by reducing the beneficial effects of adding or averaging responses, a limitation not present with units showing negative correlation (Zohary, Shadlen, and Newsome 1994). There are more unit pairs that share a positive correlation and the change in the positive correlation coefficients between NR and R trials shows a greater impact on the mean correlation. This aligns with previous evidence that changes in positive correlation specifically are the primary cause of the overall change in mean correlation. In the visual cortex, changes in orientation have been found to decrease positive correlation while negative correlation remained stable with shifts in preferred direction (Chelaru and Dragoi 2016). In a recurrent network model, a lower firing rate and decreased negative NC is generated with increased local inhibition and reduced excitation, resulting in an increased SNR (Chelaru and Dragoi 2016). However, our results lean in the direction of decreased SNR as NC increases with a decreased firing rate for our data as NR trials had both a decreased overall rate and an increased NC.

Our results show that R trials have decreased positive NC and a negative NC that most often either decreases or remains unchanged (Figure 8). The combined effect typically results in a decrease in median NC, indicating perhaps more information encoding capacity. Our results could be adhering to the trends seen in tuning direction, where the negative correlation is unchanged in response to changes in condition.

Our results support the hypothesis that NR trials should have higher NC coefficients. The lower overall correlation values during R trials aligns with previous research suggesting that decreases in correlation correspond with increased encoding capacity in similarly tuned neurons and homogeneous populations. Based on this, it is possible to conclude that NC plays a role in reward modulation in the motor cortex. Moving forward, this work has applications for the improvement of brain computer interfaces (BCI). Incorporating the context-dependent changes in NC into neural decoding models may improve decoding efficiency and increase BCI performance, as we have shown utilizing a neural critic based on classifiers of neural rate and LFP-PAC (Zhao et al. 2018; An et al. 2018). Likewise, incorporating NC into the neural state representation for a neural critic to be used in a reinforcement learning (RL) BCI should help improve the performance of such RL-BCIs, where the neural critic would track the users brain state related to reward expectation and reward prediction error and be used to autonomously update the BCI to perform better for the user (An et al. 2018; Sanchez et al. 2011; Prins, Sanchez, and Prasad 2017), or inform the decoder/policy (Zhao et al. 2018).

## Acknowledgements

We would like to thank the reviewers for their impactful input on improving this work as well as Kresimir Josic for early discussions on this topic.

## References

Abbott, L. F., and P. Dayan. 1999. ‘The effect of correlated variability on the accuracy of a population code’, Neural Comput, 11: 91–101.

An, J., T. Yadav, M. B. Ahmadi, V. S. A. Tarigoppula, and J. T. Francis. 2018. ‘Near Perfect Neural Critic from Motor Cortical Activity Toward an Autonomously Updating Brain Machine Interface’, Conf Proc IEEE Eng Med Biol Soc, 2018: 73–76.

An, J., T. Yadav, J. P. Hessburg, and J. T. Francis. 2019. ‘Reward Expectation Modulates Local Field Potentials, Spiking Activity and Spike-Field Coherence in the Primary Motor Cortex’, eNeuro, June 6 2019.

An, Junmo, Taruna Yadav, Mohammad Badri Ahmadi, Venkata Aditya Tarigoppula, and Joseph Thachil Francis. 2018. ‘Near Perfect Neural Critic from Motor Cortical Activity Toward an Autonomously Updating Brain Machine Interface’, bioRxiv.

An, Junmo, Taruna Yadav, John P Hessburg, and Joseph Thachil Francis. 2018. ‘Reward Modulates Local Field Potentials, Spiking Activity and Spike-Field Coherence in the Primary Motor Cortex’, bioRxiv.

Averbeck, B. B., and D. Lee. 2006. ‘Effects of noise correlations on information encoding and decoding’, J Neurophysiol, 95: 3633–44.

Bair, W., E. Zohary, and W. T. Newsome. 2001. ‘Correlated firing in macaque visual area MT: time scales and relationship to behavior’, J Neurosci, 21: 1676–97.

Bakhurin, K. I., V. Mac, P. Golshani, and S. C. Masmanidis. 2016. ‘Temporal correlations among functionally specialized striatal neural ensembles in reward-conditioned mice’, J Neurophysiol, 115: 1521–32.

Barreiro, Andrea K., and Cheng Ly. 2017. ‘When do correlations increase with firing rates in recurrent networks?’, PLOS Computational Biology, 13: e1005506.

Campos, M., B. Breznen, K. Bernheim, and R. A. Andersen. 2005. ‘Supplementary motor area encodes reward expectancy in eye-movement tasks’, J Neurophysiol, 94: 1325–35.

Chelaru, M. I., and V. Dragoi. 2008. ‘Efficient coding in heterogeneous neuronal populations’, Proc Natl Acad Sci U S A, 105: 16344–9.

Chelaru, M.I., and Dragoi, V. 2016. ‘Negative Correlations in Visual Cortical Networks’, Cereb Cortex, 26: 246–56.

Churchland, M. M., J. P. Cunningham, M. T. Kaufman, J. D. Foster, P. Nuyujukian, S. I. Ryu, and K. V. Shenoy. 2012. ‘Neural population dynamics during reaching’, Nature, 487: 51–6.

Churchland, M. M., B. M. Yu, J. P. Cunningham, L. P. Sugrue, M. R. Cohen, G. S. Corrado, W. T. Newsome, A. M. Clark, P. Hosseini, B. B. Scott, D. C. Bradley, M. A. Smith, A. Kohn, J. A. Movshon, K. M. Armstrong, T. Moore, S. W. Chang, L. H. Snyder, S. G. Lisberger, N. J. Priebe, I. M. Finn, D. Ferster, S. I. Ryu, G. Santhanam, M. Sahani, and K. V. Shenoy. 2010. ‘Stimulus onset quenches neural variability: a widespread cortical phenomenon’, Nat Neurosci, 13: 369–78.

Cohen, M. R., and A. Kohn. 2011. ‘Measuring and interpreting neuronal correlations’, Nat Neurosci, 14: 811–9.

Cohen, M. R., and J. H. Maunsell. 2009. ‘Attention improves performance primarily by reducing interneuronal correlations’, Nat Neurosci, 12: 1594–600.

Cohen, M. R., and W. T. Newsome. 2008. ‘Context-dependent changes in functional circuitry in visual area MT’, Neuron, 60: 162–73.

de la Rocha, J., B. Doiron, E. Shea-Brown, K. Josic, and A. Reyes. 2007. ‘Correlation between neural spike trains increases with firing rate’, Nature, 448: 802–6.

Dura-Bernal, S., X. L. Zhou, S. A. Neymotin, A. Przekwas, J. T. Francis, and W. W. Lytton. 2015. ‘Cortical Spiking Network Interfaced with Virtual Musculoskeletal Arm and Robotic Arm’, Frontiers in Neurorobotics, 9.

Dushanova, J., and J. Donoghue. 2010. ‘Neurons in primary motor cortex engaged during action observation’, Eur J Neurosci, 31: 386–98.

Ecker, A. S., P. Berens, G. A. Keliris, M. Bethge, N. K. Logothetis, and A. S. Tolias. 2010. ‘Decorrelated neuronal firing in cortical microcircuits’, Science, 327: 584–7.

Francis, J. T., and W. Song. 2011. ‘Neuroplasticity of the sensorimotor cortex during learning’, Neural Plast, 2011: 310737.

Galan, R. F., N. Fourcaud-Trocme, G. B. Ermentrout, and N. N. Urban. 2006. ‘Correlation-induced synchronization of oscillations in olfactory bulb neurons’, J Neurosci, 26: 3646–55.

Gu, Y., S. Liu, C. R. Fetsch, Y. Yang, S. Fok, A. Sunkara, G. C. DeAngelis, and D. E. Angelaki. 2011. ‘Perceptual learning reduces interneuronal correlations in macaque visual cortex’, Neuron, 71: 750–61.

Hansen, B. J., M. I. Chelaru, and V. Dragoi. 2012. ‘Correlated variability in laminar cortical circuits’, Neuron, 76: 590–602.

Herrero, J. L., M. A. Gieselmann, M. Sanayei, and A. Thiele. 2013. ‘Attention-induced variance and noise-correlationreduction in macaque V1 is mediated by NMDA receptors’, Neuron, 78: 729–39.

Hertz, J. 2010. ‘Cross-correlations in high-conductance states of a model cortical network’, Neural Comput, 22: 427–47.

Hessburg, J. P., Y. Zhao, and J. T. Francis. 2019. ‘Divisive Normalization by Valence and Motivational Intensity in the Primary Sensorimotor Cortices (PMd, M1, S1)’, bioRxiv.

Jeanne, J. M., T. O. Sharpee, and T. Q. Gentner. 2013. ‘Associative learning enhances population coding by inverting interneuronal correlation patterns’, Neuron, 78: 352–63.

Kohn, A., and M. A. Smith. 2005. ‘Stimulus dependence of neuronal correlation in primary visual cortex of the macaque’, J Neurosci, 25: 3661–73.

Komiyama, T., T. R. Sato, D. H. O’Connor, Y. X. Zhang, D. Huber, B. M. Hooks, M. Gabitto, and K. Svoboda. 2010. ‘Learning-related fine-scale specificity imaged in motor cortex circuits of behaving mice’, Nature, 464: 1182–6.

Kunori, N., R. Kajiwara, and I. Takashima. 2014. ‘Voltage-sensitive dye imaging of primary motor cortex activity produced by ventral tegmental area stimulation’, J Neurosci, 34: 8894–903.

Lee, J., and J. H. Maunsell. 2009. ‘A normalization model of attentional modulation of single unit responses’, PLoS One, 4: e4651.

Louie, K., L. E. Grattan, and P. W. Glimcher. 2011. ‘Reward value-based gain control: divisive normalization in parietal cortex’, J Neurosci, 31: 10627–39.

Marsh, B. T., V. S. Tarigoppula, C. Chen, and J. T. Francis. 2015. ‘Toward an autonomous brain machine interface: integrating sensorimotor reward modulation and reinforcement learning’, J Neurosci, 35: 7374–87.

McNiel, D. B., M. Bataineh, J. S. Choi, J.P. Hessburg, and J. T. Francis. 2016. ‘Classifier Performance in Primary Somatosensory Cortex Towards Implementation of a Reinforcement Learning Based Brain Machine Interface’, IEEE Southern Biomedical Engineering Conference 2016.

McNiel, D. B., J. S. Choi, J.P. Hessburg, and J. T. Francis. 2016a. ‘Reward value is encoded in primary somatosensory cortex and can be decoded from neural activity during performance of a psychophysical task’, IEEE EMBS 2016 Conference proceding (Aug 2016).

McNiel, David B, John S Choi, John P Hessburg, and Joseph T Francis. 2016b. “Reward value is encoded in primary somatosensory cortex and can be decoded from neural activity during performance of a psychophysical task.” In Engineering in Medicine and Biology Society (EMBC), 2016 IEEE 38th Annual International Conference of the, 3064–67. IEEE.

Mitchell, J. F., K. A. Sundberg, and J. H. Reynolds. 2007. ‘Differential attention-dependent response modulation across cell classes in macaque visual area V4’, Neuron, 55: 131–41.

Mitchell, J. F., Sundberg, K. A. and Reynolds, J. H. 2009. ‘Spatial attention decorrelates intrinsic activity fluctuations in macaque area V4’, Neuron, 63: 879–88.

Molina-Luna, K., A. Pekanovic, S. Rohrich, B. Hertler, M. Schubring-Giese, M. S. Rioult-Pedotti, and A. R. Luft. 2009. ‘Dopamine in motor cortex is necessary for skill learning and synaptic plasticity’, PLoS One, 4: e7082.

Muller, M. M., W. Teder-Salejarvi, and S. A. Hillyard. 1998. ‘The time course of cortical facilitation during cued shifts of spatial attention’, Nat Neurosci, 1: 631–4.

Musallam, S., B. D. Corneil, B. Greger, H. Scherberger, and R. A. Andersen. 2004. ‘Cognitive control signals for neural prosthetics’, Science, 305: 258–62.

Nirenberg, S., and P. E. Latham. 2003. ‘Decoding neuronal spike trains: how important are correlations?’, Proc Natl Acad Sci U S A, 100: 7348–53.

Okun, M., N. Steinmetz, L. Cossell, M. F. Iacaruso, H. Ko, P. Bartho, T. Moore, S. B. Hofer, T. D. Mrsic-Flogel, M. Carandini, and K. D. Harris. 2015. ‘Diverse coupling of neurons to populations in sensory cortex’, Nature, 521: 511–15.

Padmanabhan, K., and N. N. Urban. 2010. ‘Intrinsic biophysical diversity decorrelates neuronal firing while increasing information content’, Nat Neurosci, 13: 1276–82.

Platt, M. L., and P. W. Glimcher. 1999. ‘Neural correlates of decision variables in parietal cortex’, Nature, 400: 233–8.

Ramakrishnan, A., Y. W. Byun, K. Rand, C. E. Pedersen, M. A. Lebedev, and M. A. L. Nicolelis. 2017. ‘Cortical neurons multiplex reward-related signals along with sensory and motor information’, Proc Natl Acad Sci U S A, 114: E4841–E50.

Ramkumar, P., B. Dekleva, S. Cooler, L. Miller, and K. Kording. 2016. ‘Premotor and Motor Cortices Encode Reward’, PLoS One, 11: e0160851.

Reid, R. C., and J. M. Alonso. 1995. ‘Specificity of monosynaptic connections from thalamus to visual cortex’, Nature, 378: 281–4.

Renart, A., J. de la Rocha, P. Bartho, L. Hollender, N. Parga, A. Reyes, and K. D. Harris. 2010. ‘The asynchronous state in cortical circuits’, Science, 327: 587–90.

Romo, R., A. Hernandez, A. Zainos, and E. Salinas. 2003. ‘Correlated neuronal discharges that increase coding efficiency during perceptual discrimination’, Neuron, 38: 649–57.

Ruff, D. A., J. J. Alberts, and M. R. Cohen. 2016. ‘Relating normalization to neuronal populations across cortical areas’, J Neurophysiol, 116: 1375–86.

Ruff, D. A., and M. R. Cohen. 2014. ‘Global cognitive factors modulate correlated response variability between V4 neurons’, J Neurosci, 34: 16408–16.

Schultz, W., P. Dayan, and P. R. Montague. 1997. ‘A neural substrate of prediction and reward’, Science, 275: 1593–9.

Shadlen, M. N., and W. T. Newsome. 1998. ‘The variable discharge of cortical neurons: implications for connectivity, computation, and information coding’, J Neurosci, 18: 3870–96.

Shuler, M. G., and M. F. Bear. 2006. ‘Reward timing in the primary visual cortex’, Science, 311: 1606–9.

Smith, M. A., X. Jia, A. Zandvakili, and A. Kohn. 2013. ‘Laminar dependence of neuronal correlations in visual cortex’, J Neurophysiol, 109: 940–7.

Tanaka, S. C., K. Doya, G. Okada, K. Ueda, Y. Okamoto, and S. Yamawaki. 2004. ‘Prediction of immediate and future rewards differentially recruits cortico-basal ganglia loops’, Nat Neurosci, 7: 887–93.

Tarigoppula, A., J. S. Choi, J.P. Hessburg, D. B. McNiel, B. T. Marsh, and J..T. Francis. 2018a. ‘Motor Cortex Encodes A Value Function Consistent With Reinforcment Learning’, bioRxiv.

Tarigoppula, Venkata S Aditya, John S Choi, John H Hessburg, David B McNiel, Brandi T Marsh, and Joseph Thachil Francis. 2018b. ‘Primary Motor Cortex Encodes A Temporal Difference Reinforcement Learning Process’, bioRxiv.

Tkach, D., J. Reimer, and N. G. Hatsopoulos. 2007. ‘Congruent activity during action and action observation in motor cortex’, J Neurosci, 27: 13241–50.

Toda, K., Y. Sugase-Miyamoto, T. Mizuhiki, K. Inaba, B. J. Richmond, and M. Shidara. 2012. ‘Differential encoding of factors influencing predicted reward value in monkey rostral anterior cingulate cortex’, PLoS One, 7: e30190.

Tolhurst, D. J., J. A. Movshon, and A. F. Dean. 1983. ‘The statistical reliability of signals in single neurons in cat and monkey visual cortex’, Vision Res, 23: 775–85.

Tremblay, L., and W. Schultz. 1999. ‘Relative reward preference in primate orbitofrontal cortex’, Nature, 398: 704–8.

Vaadia, E., I. Haalman, M. Abeles, H. Bergman, Y. Prut, H. Slovin, and A. Aertsen. 1995. ‘Dynamics of neuronal interactions in monkey cortex in relation to behavioural events’, Nature, 373: 515–8.

Wilson, C. J. 2013. ‘Active decorrelation in the basal ganglia’, Neuroscience, 250: 467–82.

Zhao, Y., J. P. Hessburg, J. N. A. Kumar, and J. T. Francis. 2018. ‘Paradigm Shift in Sensorimotor Control Research and Brain Machine Interface Control: The Influence of Context on Sensorimotor Representations’, Frontiers in Neuroscience, 12.

Zohary, E., M. N. Shadlen, and W. T. Newsome. 1994. ‘Correlated neuronal discharge rate and its implications for psychophysical performance’, Nature, 370: 140–3.

